# Contributions of Folded and Disordered Domains to RNA Binding by HNRNPR

**DOI:** 10.1101/2025.05.01.651718

**Authors:** Bryan B. Guzmán, Grant A. Goda, Alli Jimenez, Justin G. Martyr, Yue Hu, Francisco F. Cavazos, Maria M. Aleman, Daniel Dominguez

## Abstract

RNA binding proteins (RBPs) interact with and tightly regulate the fate of messenger RNAs but how RNA targets are recognized remains a challenging question. RBPs often contain multiple domains known to directly bind RNA, such as RNA recognition motifs (RRMs), as well as domains whose RNA binding capacity remains incompletely understood, *e.g*., low complexity domains (LCDs). Here, we dissect HNRNPR, an RBP with three RRMs and an arginine-glycine rich (RG-rich) LCD. We apply unbiased high-throughput biochemical approaches and identify critical RNA binding domains that confer specificity. We show that not all RRMs contribute equally to binding and find that RRM3, along with a downstream C-terminal charged region, are required for RNA binding. We find that HNRNPR also binds RNA G-quadruplexes (rG4s) and map multiple rG4 binding sites including RRM3 with the C-terminal charged region and RG-rich regions within the LCD. We dissect rG4 specificity for the full length HNRNPR and LCD using a newly created RNA pool focused on rG4s and reveal that binding is dependent on RNA folding and find specific rG4 features that enhance HNRNPR-rG4 interactions. Our work highlights the complexity of RBP-RNA interactions and motivates the study of disordered regions as RNA binding domains.

## INTRODUCTION

Regulation of mRNA biogenesis and fate is central to gene expression control. RNA binding proteins (RBPs) directly and specifically interact with RNA to modulate splicing, localization, and translation (1–4). RBPs also complex with non-coding RNAs (*e.g*., *XIST* and *7SK*) to exert transcriptional control (5–7). Thus, a mechanistic understanding of how RBPs select their RNA targets is essential for understanding gene regulation.

The best understood RBPs possess one or more canonical RNA binding domains (RBD), such as the RNA recognition motif (RRM), HNRNPK-homology domain (KH), or zinc finger domain (ZNF) (8). Canonical RBDs are generally globular and adopt stereotypical folds that enable specific RNA binding [reviewed in (4)]. Despite dissection of RBDs over past decades, structural and biochemical studies continue to uncover unexpected binding features that have made it a challenge to infer RNA specificity from amino acid composition (9, 10). Furthermore, many RBPs contain multiple RBDs thought to work cooperatively to enhance sequence specificity and affinity (11, 12); yet, in many cases the contributions of each RBD to target selection remains incompletely understood.

Unbiased methodologies continue to identify new proteins and/or domains with RNA binding capacity (13, 14). Of interest to this study are intrinsically disordered domains, especially those with skewed amino acid content, such as those enriched in arginine and glycine (RG-rich) or serine and arginine (SR-rich) [reviewed in (15) and (16)]. These domains, also termed low-complexity domains (LCDs), are enriched among RBPs (17) and have been shown to interact with RNA in sequence-specific and sequence-independent manners (18, 19). For example, the RG-rich disordered LCD of FMRP interacts with guanine-rich RNA motifs that can fold into RNA G-quadruplexes (rG4s), a regulatory element important for mRNA localization to neuronal projections (20, 21). In a separate structural study, the disordered RG-rich domain of FUS was shown to primarily interact with RNA in a sequence-independent manner and primarily stabilizes interactions further mediated by the FUS ZNF and RRM (22). Other RG-rich LCDs, primarily found in RBPs, have also been shown to bind RNA with varying affinities and specificities, with the rG4 fold emerging as a recurrent LCD binding partner [reviewed in (23)].

Despite decades of research on the HNRNP family of RBPs, detailed RNA-binding modes and their functional consequences for some family members remain unclear. HNRNPR is a multifaceted RBP with regulatory activities in transcription, splicing, localization, and translation (24–27). Phenotypically, loss of HNRNPR disrupts axonal growth in motor neurons (28), alters the DNA damage response (29), and modulates global transcription (24). One study found that HNRNPR binds 3’ untranslated regions (UTRs) of target mRNAs to mediate mRNA localization (26). A key example is the interaction of HNRNPR with the 3’ UTR of beta-actin (*ACTB*) mRNA that, along with the SMN protein, mediates axonal translocation of said mRNA (28, 30). HNRNPR-RNA interactome studies have revealed the non-coding *7SK* RNA as a primary binding partner (26), but the binding mechanism remains unclear. While HNRNPR-*7SK* interactions are thought to control transcription by remodeling the *7SK* ribonucleoprotein complex (RNP), intriguingly, an HNRNPR-*7SK* RNP has also been proposed to traffic mRNAs to neuronal projections where local protein synthesis takes place (26).

Mutations in HNRNPR are linked to human disease (31, 32). Dujikers et al., identified HNRNPR mutations in individuals with multi-system abnormalities impacting bone, brain, and the reproductive system (32). In addition, Gillentine et al. recently reported a set of deleterious mutations in *HNRNP* genes, including *HNRNPR* (31). Both studies have primarily found disease-related point mutants and truncations impacting the disordered LCD of HNRNPR, indicating this region is of functional importance.

Here we dissect the RNA selectivity of HNRNPR using unbiased high-throughput biochemical approaches. We define the critical HNRNPR RBDs required for binding to AU-rich RNAs and identify the rG4 structural motif as a secondary target. We further evaluate the rG4 specificity of full length HNRNPR and the LCD alone using a diverse pool of thousands of rG4s with varying sequence features. We show that binding of the LCD to rG4s required the folded rG4 structure and highlight preferences for specific rG4s dependent on loop length and nucleotide composition. Unexpectedly, we found that HNRNPR RRMs require a proximal charged region outside of RRM3 to confer high-affinity RNA interactions to both AU-rich and rG4 RNA, and further demonstrate that AU-rich RNA and rG4s competed for HNRNPR binding. Our studies also highlight the multi-valency of HNRNPR-rG4 interactions through mapping multiple rG4 binding sites within the LCD and show that arginine-content within the LCD was critical for high-affinity interactions. This in-depth biochemical study generates novel insights into the complex RNA recognition of HNRNPR through its RRMs and LCD, providing a unique perspective that may be widely applicable to other RBPs and their specific interactions with target RNAs.

## RESULTS

### The RNA specificity of HNRNPR

HNRNPR is a prototypical RBP with three RRMs and a disordered domain rich in arginine and glycine (**Fig. 1A**). HNRNPR RNA targets have been identified (*e.g*., *7SK* and *ACTB*) by several methodologies but little is known about the mechanism driving HNRNPR-RNA recognition. We set out to answer this question, with the goal of identifying the critical domains and their associated sequence specificities. We purified recombinant full length (FL) HNRNPR (residues 1-636) and performed RNA Bind-n-Seq (RBNS), a high-throughput *in vitro* binding assay that discovers RNA specificity in an unbiased fashion by allowing a protein to select binding partners from an extremely diverse set of randomized RNAs (33, 34) (**Fig. 1B**).

**Figure 1.**
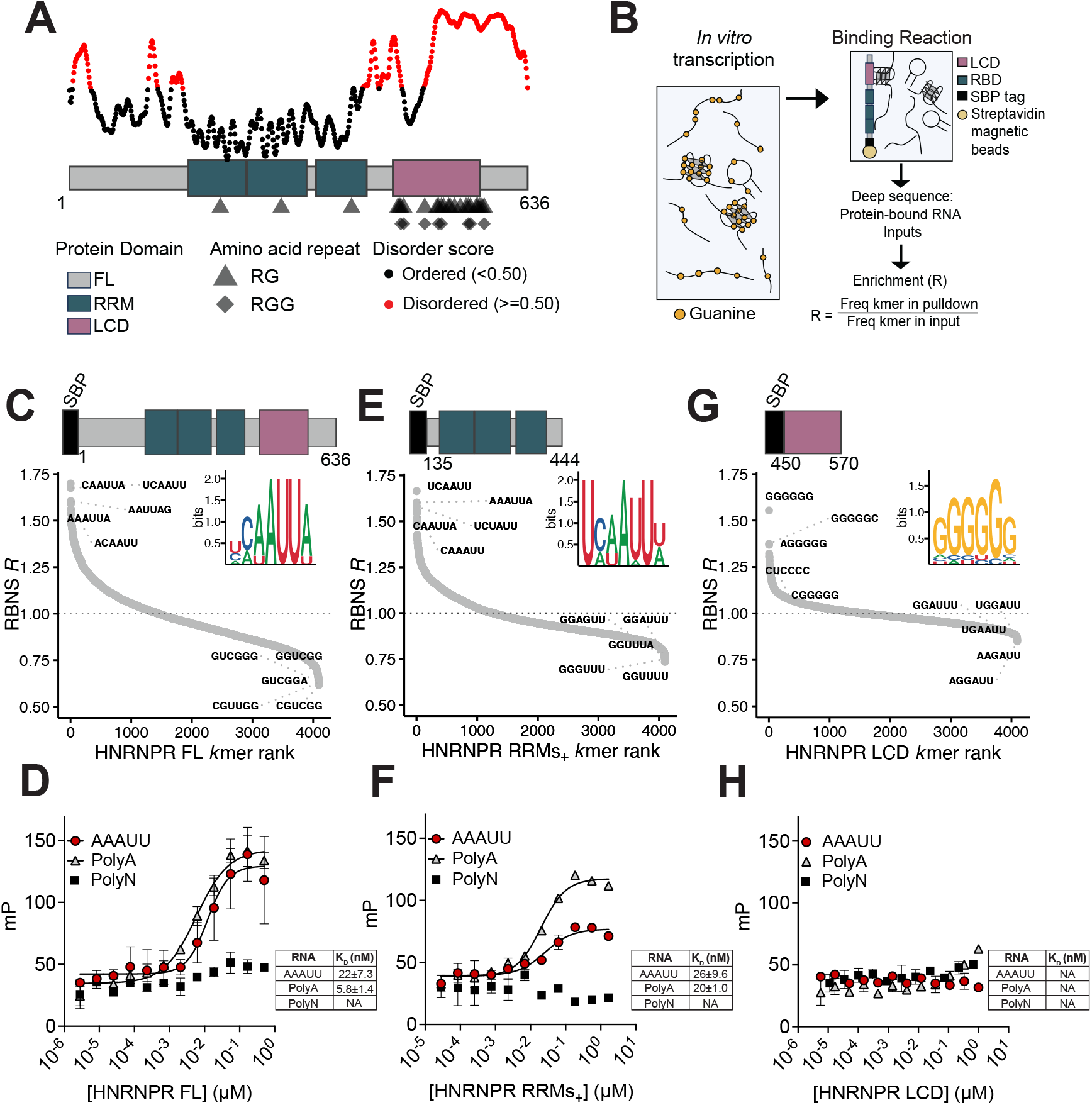
HNRNPR RNA binding specificity. A) Bottom, schematic of HNRNPR and its respective domains: Full length (FL) (grey), RRMs (dark teal) and LCD (light purple). Below the schematic, RG (triangle) or RGG (diamond) repeats in HNRNPR are plotted. Top, disorder propensity plot (IUPRED2) of HNRNPR with ordered amino acids (black dot) and disordered amino acids (red dot) indicated. B) Schematic of RNA Bind-n-Seq (RBNS) workflow. C) Top, schematic of protein fragment used for RBNS. Bottom, ranked scatter plots of RBNS enrichments of all 6mers for HNRNPR FL. D) Fluorescence polarization (FP) binding curves (N=3) for HNRNPR FL incubated with AAAUU RNA (red dot), polyA RNA (grey triangle) and polyN RNA (black square). Data are mean ± standard deviation (SD). E) Top, schematic of protein fragment used for RBNS. Bottom, ranked scatter plots of RBNS enrichments of all 6mers for HNRNPR RRMs_+_. F) FP binding curves (N=3) for HNRNPR RRMs_+_ incubated with AAAUU RNA (red dot), polyA RNA (grey triangle) and polyN RNA (black square). Data are mean ± SD. G) Top, schematic of protein fragment used for RBNS. Bottom, ranked scatter plots of RBNS enrichments of all 6mers for HNRNPR LCD. H) FP binding curves (N=3) for HNRNPR LCD incubated with AAAUU RNA (red dot), polyA RNA (grey triangle) and polyN RNA (black square). Data are mean ± SD.

RBNS on FL HNRNPR revealed enrichments for AU-rich motifs (**Fig. 1C**). In addition to AU-rich preferences, the most enriched sequences also contained a cytosine within the motif, generally preceding an adenine (*e.g*., CAAUUA). *K*mer enrichments across replicates and different protein concentrations were reproducible and highly correlated (**Fig. S1A, S1D**). RBNS data also enables determination of RNA structure preferences of motifs bound by a given protein (33). *In silico* RNA folding analysis of millions of bound RNAs showed that, like most RRM-containing RBPs, HNRNPR preferred motifs in a single stranded context (**Fig. S1J**). CAAUUA sequences were unstructured (low base pair probabilities), and generally, the most enriched motifs were the most unstructured (**Fig. S1J**).

RBNS was also carried out on a fragment containing all three RRMs plus 30 flanking amino acids N- and C-terminal (residues 135-444, RRMs_+_). RRMs_+_ preferred nearly identical motifs as the FL protein, with *k*mer enrichments showing a high degree of correlation between FL and RRMs_+_ (**Fig. 1E, Fig. S1G**). As expected, the RRMs_+_ had similar preferences for unstructured motifs (**Fig. S1K**).

To determine the affinity of HNRNPR FL and RRMs_+_ towards AU-rich RNAs, we measured the interaction using an orthogonal approach. We designed multiple AU-rich fluorescein-labeled RNA oligos and tested their binding to FL protein and RRMs_+_ by fluorescence polarization (FP) (**Fig. 1D, 1F**). Both FL and RRMs_+_ displayed high affinity towards these sequences. Furthermore, neither FL nor RRMs_+_ alone interacted with a randomized RNA sequence (**Fig. 1D, 1F**), confirming sequence specificity. Together, these data demonstrate that FL HNRNPR, via the RRM domains, bound with nanomolar affinity to single-stranded AU-rich motifs, with a preference towards a cytosine prefix (*e.g*., CAAUUA).

### HNRNPR RG-rich LCD interacts with rG4s

While RRMs are well-established drivers of RNA binding, emerging data indicate that low complexity domains (LCDs) within RBPs, particularly those with RG-rich or RGG repeats, also directly interact with RNA or enhance RRM RNA binding affinity [(22) and reviewed in (19)]. RBNS on the HNRNPR LCD alone (residues 450-570) uncovered a clear preference towards G-rich motifs (**Fig. 1G**). This specificity was in stark contrast to the top AU-rich RNA motifs bound by HNRNPR FL and RRMs_+_. *K*mer enrichments for RRMs_+_ vs LCD were markedly different and of the top 5 *k*mers bound by RRMs_+_ or the LCD, none overlapped (**Fig. 1E, 1G, Fig. S1F**). FP experiments confirmed the LCD did not interact with AU-rich RNA with appreciable affinity, nor did it interact with a randomized RNA sequence (**Fig. 1H**). Thus, HNRNPR contains multiple domains capable of binding RNA with significantly different specificities.

Previous work on RG-rich disordered regions within RBPs has shown that they interact with G-rich RNAs that fold into RNA G-quadruplexes (rG4s) (20, 21, 35, 36). rG4s form from a core motif of (G_2-5_N_1-7_)_≥4_ by Hoogsteen base pairing interactions which stabilize four guanine residues, one from each polyG sequence (G-tract), that make up a planar G-tetrad (**Fig. 2A**). rG4 sequences are over-represented in the transcriptome, particularly in untranslated regions, and have been shown to function as regulatory elements for splicing, translation, and localization (37–39).

**Figure 2.**
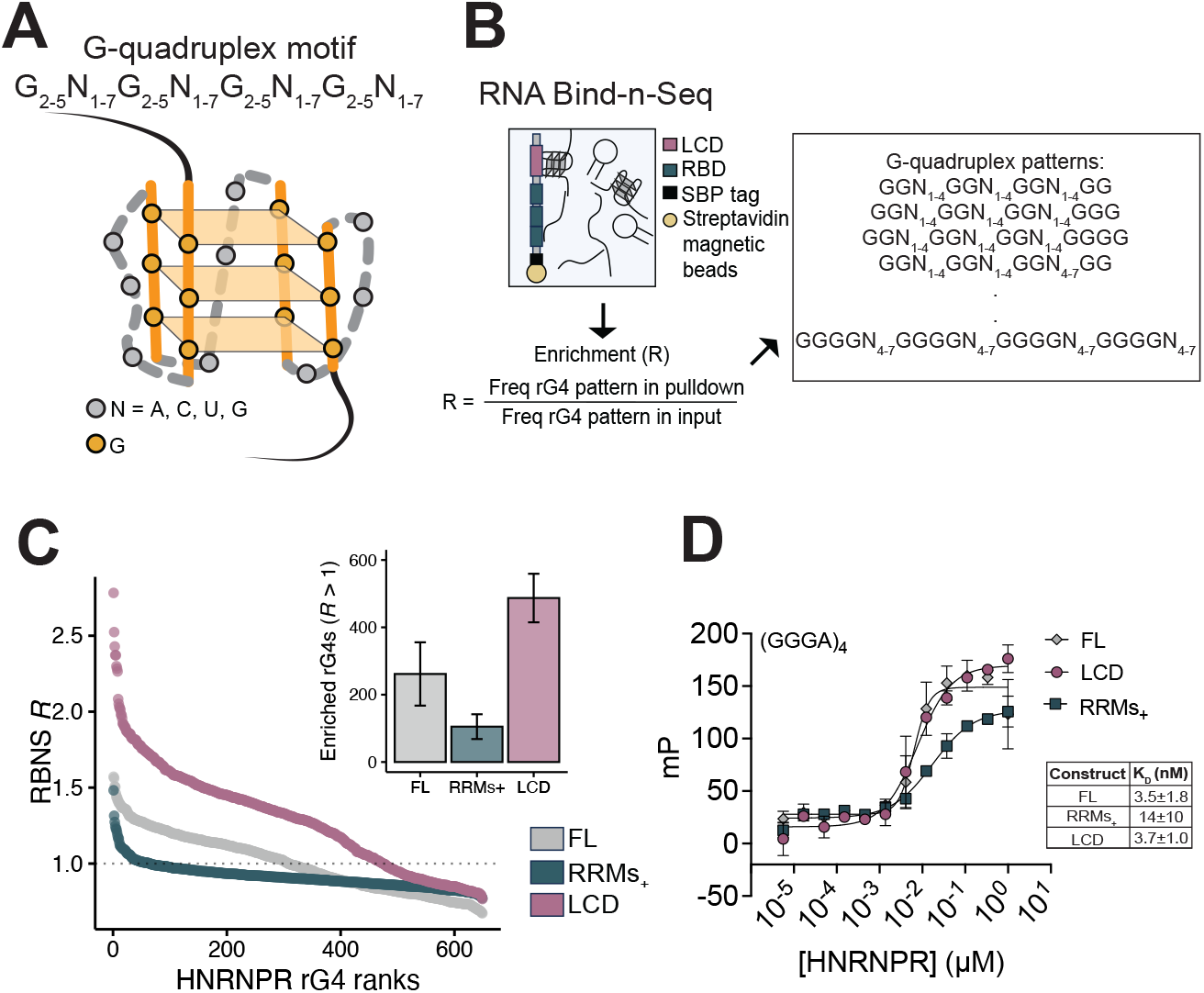
HNRNPR binds RNA G-quadruplexes. A) Schematic of an RNA G-quadruplex (rG4). B) Schematic of RBNS workflow with a modification: enrichments are calculated based on a curated list of 649 rG4 patterns. C) Ranked scatter plot for enrichments of rG4 patterns determined by RBNS for: FL (grey), RRMs_+_ (dark teal) and LCD (light purple). Inset, number of rG4 patterns that have an enrichment value above 1 by RBNS for FL, RRMs_+_ and LCD. Data are mean ± standard deviation (SD). D) FP binding curves (N=3) for FL, RRMs_+_ and LCD incubated with (GGGA)_4_ RNA. Data are mean ± SD.

Due to the G-rich specificity displayed by the LCD, we set out to determine if FL HNRNPR interacts with rG4s. We harnessed the depth of our RBNS data to search for long motifs that are relatively rare, such as rG4s (**Fig. 2B**). For example, there are ∼65,000 8mers, the shortest possible length of a canonical rG4, and quantifying each of these sequences with confidence requires a significant number of protein-associated reads. While rG4s have a core fold, they can have varying topologies and stabilities depending on their sequence composition. We curated a list of rG4 sub-sequences from the general rG4 motif (G_2-5_N_1-7_) _≥4_. These putative rG4 sequences vary in the number of Gs in each G-tract and in the number of nucleotides per loop (either 1-4 or 4-7 nts in length) (**Table S1**). We calculated the enrichments for these rG4 patterns by assessing how frequently they appeared in protein associated reads vs input reads (**Fig. 2B**). Thus, an enrichment value above 1 suggests that the recombinant protein or domain associates with that specific pattern. Using this metric, we found that FL does have a degree of enrichment for rG4-containing sequences (**Fig. 2C**). Both HNRNPR fragments, the LCD and RRMs_+_ alone, enriched for rG4-forming sequences, although RRMs_+_ bound to a lesser extent (**Fig. 2C**, discussed below). Of note, the LCD displayed significantly higher enrichments for rG4s than the FL protein (**Fig. 2C**). *K*mer enrichments across replicates were reproducible and highly correlated (**Fig. S2A-C**). HNRNPR-rG4 interactions were validated via FP using an rG4-forming RNA oligo, where HNRNPR FL and LCD bound with nanomolar affinity (**Fig. 2D**). To further confirm HNRNPR specificity towards rG4s, we performed a competition assay using FP. In this experiment, we incubated protein with fluorescein labeled rG4 (GGGA)_4_ then titrated in unlabeled rG4 (GGGA)_4_. Indeed, unlabeled rG4 was able to outcompete the fluorescein labeled rG4 (**Fig. S2D**). These data strongly indicate that HNRNPR FL interacts with rG4s; however, it raises the question of whether FL and LCD have specificity towards different rG4 sequences.

### HNRNPR specificity towards rG4s

As previously mentioned, the canonical rG4-forming pattern allows for significant sequence variability, which complicates understanding general RBP specificity towards rG4s. To address rG4 specificity of HNRNPR beyond the core rG4 motif pattern, we took advantage of a structural rG4 (stG4) pool we recently designed (40). This curated stG4 pool was created with the goal of understanding sequence elements that dictate rG4 stability as well as molecular recognition, including protein binding. Briefly, the stG4 pool was designed through combinations of G-tracts, loop composition, and loop length, all potential determinants of RBP-rG4 specificity (**Fig. 3A**). rG4 formation within the pool can further be promoted or inhibited by *in vitro* transcribing RNA with either guanine or with 7-deaza-guanine (7dG), which can hinder Hoogsteen base pairing due to the substitution of a critical nitrogen (N7) to a carbon (41, 42). In a related study, we used this stG4 pool to carry out reverse transcription (RT) stop assays to evaluate sequences capable of folding into rG4s for thousands of RNAs in parallel (40). These experiments resulted in an RT Stop Score, where a score of -2 or lower indicates a ‘folded rG4’ (Methods detailed elsewhere (40)). With this specialized rG4 pool, we set out to understand HNRNPR FL and LCD specificity towards rG4s and the impact of the fold on binding.

**Figure 3.**
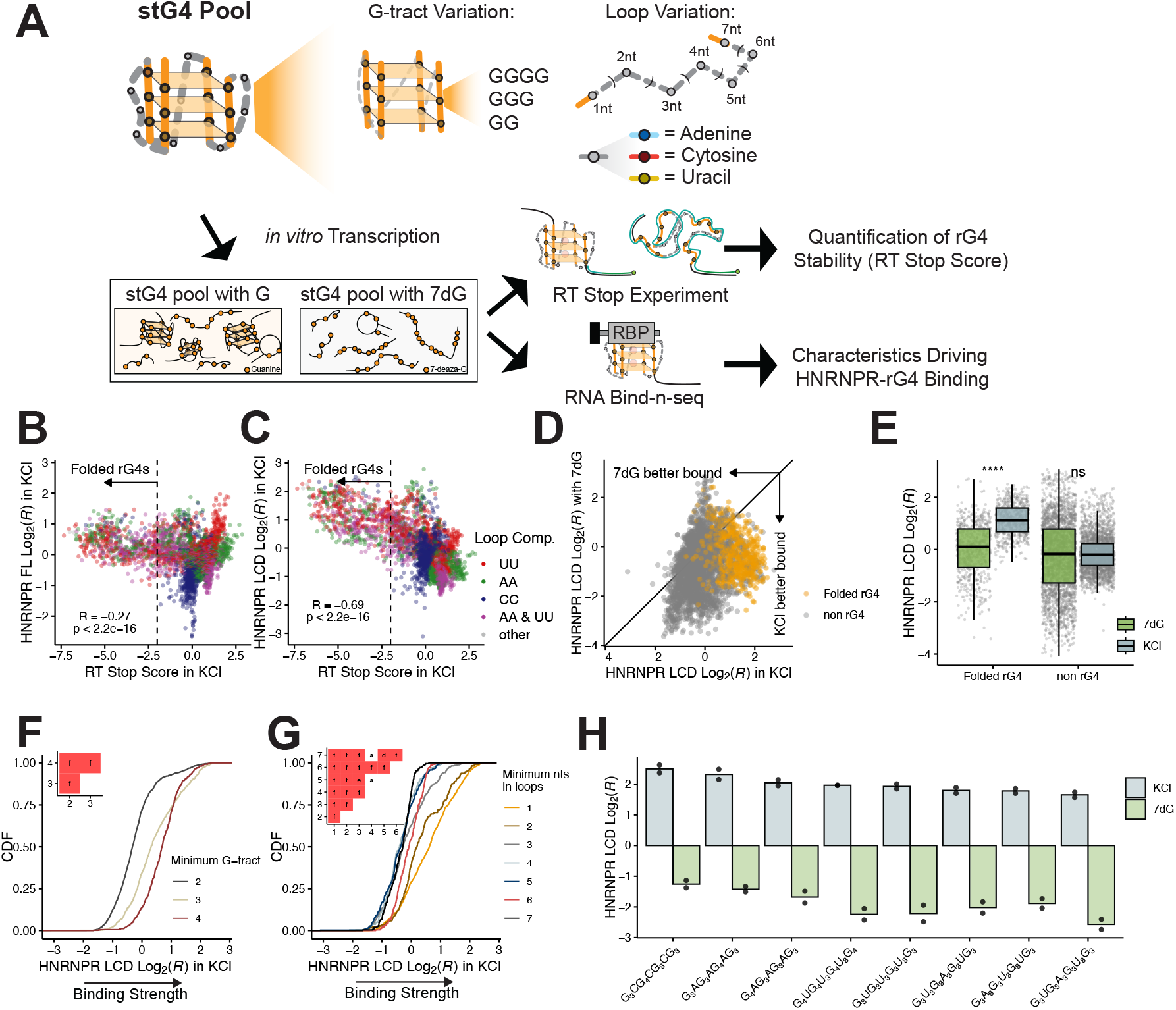
HNRNPR specificity towards rG4s. A) Schematic of the stG4 pool with breakdown of different RNA sequences within the pool and applications of the pool adapted from (40). Comparison of log_2_ RBNS enrichment for B) FL and C) LCD with guanine in KCl (y-axis) versus RT Stop Score of rG4s (x-axis), where an RT Stop Score of < -2 denotes folded rG4s. Colors represent the nucleotide composition in the rG4 loops. The minimum requirement for classification was to have the same nucleotide twice in adjacent positions or consecutives loops. D) Log_2_ RBNS enrichment for LCD with 7dG rG4s (y-axis) versus log_2_ RBNS enrichment for LCD with guanine rG4s in KCl (x-axis). Grey points represent non rG4s (RT Stop Score above -2) and gold points represent folded rG4s (RT Stop Score below -2). E) Boxplot shows the log_2_ RBNS enrichment for LCD for non rG4 and folded rG4 with 7dG RNA (light green) and with guanine in KCl (light blue). Significance was determined by Wilcoxon test. Significance marks are as follows: **** (p ≤ 0.0001), ns (not significant). F) Cumulative distribution function (CDF) of log_2_ RBNS enrichment for the LCD with guanine in KCl separated by the minimum G-tract allowed in an oligo. Inset shows p-values determined by a two-sided KS test corrected via the BH procedure. Red denotes significance and values are as follows: f (p≤0.0001). G) CDF of log_2_ RBNS enrichment for LCD with guanine in KCl separated by minimum nucleotide counts allowed in the rG4 loops in an oligo. Inset shows p-values determined by a two-sided KS test corrected via the BH procedure. Red denotes significance and values are as follows: a (ns), d (p≤0.01), e (p≤0.001), f (p≤0.0001). H) Log_2_ RBNS enrichment (y-axis) of specific examples of oligos bound by LCD with 7dG RNA (light green) and guanine in KCl (light blue). Bars represent the mean of enrichment and points denote individual enrichments.

We performed a massively parallel binding assay between this stG4 pool and HNRNPR FL as well as the LCD. Binding enrichments were determined between HNRNPR and ∼6000 RNAs with strong reproducibility between binding replicates (**Fig. S3A-D**). We first determined the extent to which HNRNPR binding correlated with rG4 strength (RT Stop Score), with intent to understand whether the strength of the rG4 fold impacted HNRNPR binding. Binding to folded rG4s was readily apparent for both HNRNPR FL and LCD (**Fig. 3B, 3C**). In addition, a population of RNAs predicted to be unfolded were also bound by HNRNPR FL, an effect likely driven by RRM-RNA binding (**Fig. 3B**). Inspection of non-rG4 sequences bound by FL HNRNPR demonstrated that these sequences contained loops with polyA and polyU motifs, consistent with RRM-mediated interactions (**Fig. 3B**). The same experiment with the LCD showed preferential binding to folded rG4 sequences and less binding to non-rG4 sequences (**Fig. 3C**). When comparing LCD versus FL, we observed a stronger correlation between rG4 folding and LCD binding than what we observed for FL HNRNPR (**Fig. 3B, 3C**).

To confirm that binding was mediated by the rG4 structure, we performed the same binding experiment with RNA where 7dG was substituted for guanine to preclude rG4 formation with limited alterations to the RNA sequence. Generally, this substitution led to a decrease in binding to sequences that were predicted to be strong rG4s for both the FL and LCD (**Fig. 3D, Fig. S3E**). The decrease in rG4 binding in RNAs made with 7dG was significant for both FL and LCD and only impacted RNAs capable of forming rG4s, while non-rG4 sequences were less affected by 7dG substitution (**Fig. 3E, Fig. S3F**). However, it should be noted that a small set of sequences strongly bound both FL and LCD even with 7dG substitutions (**Fig. 3D, Fig. S3E**). We expect binding by FL in these cases was mediated by AU-rich sequences within these loops and speculate that the LCD may bind and stabilize weak rG4s as has been shown in previous work (43).

We next evaluated the effect of G-tract length on HNRNPR-rG4 binding, focusing on LCD recognition. For the LCD, binding significantly increased as the number of Gs within each G-tract increased, thus GGGG>GGG>GG (**Fig. 3F**). Evaluation of loop length demonstrated that the LCD bound better to short loops (1 nt) and there was a step-wise decrease in binding strength with increasing loop length (**Fig. 3G**). Evaluation of specific rG4 sequences further highlighted the dependence of rG4 folding on binding (**Fig. 3H**). For full length protein, we noted that longer loops (6 and 7 nts) were bound better than intermediate loops of 3, 4, and 5 nucleotides (**Fig. S3J**). We interpret this as HNRNPR FL’s capability to bind both putative rG4s that may be unstructured and contain AU-rich sequences to engage the RRM. Indeed, excluding long loop rG4s with high AU-rich content in their loops showed these had virtually no enrichment for HNRNPR FL (**Fig. S3K**).

### A complex RNA binding mode by HNRNPR RRMs and a C-terminal helix

*AU-rich RNA:* We next sought to identify the minimal portions of HNRNPR that bind AU-rich as well as rG4 RNAs (discussed below). We generated a minimal recombinant protein containing exclusively the RRMs as annotated by UniProt (residues 165-411, minRRMs), but to our surprise found no detectable AU-rich RNA binding with this fragment (**Fig. 4A**). Thus, the three RRMs in their minimal form are insufficient to confer high-affinity RNA interactions. However, as shown above, a version of the protein with ∼30 additional amino acids flanking N- and C-terminal to the RRMs (RRMs_+_, residues 135-444), bound AU-rich RNAs (**Fig. 1E, 1F**), indicating that these flanking regions are critical for RNA binding.

**Figure 4.**
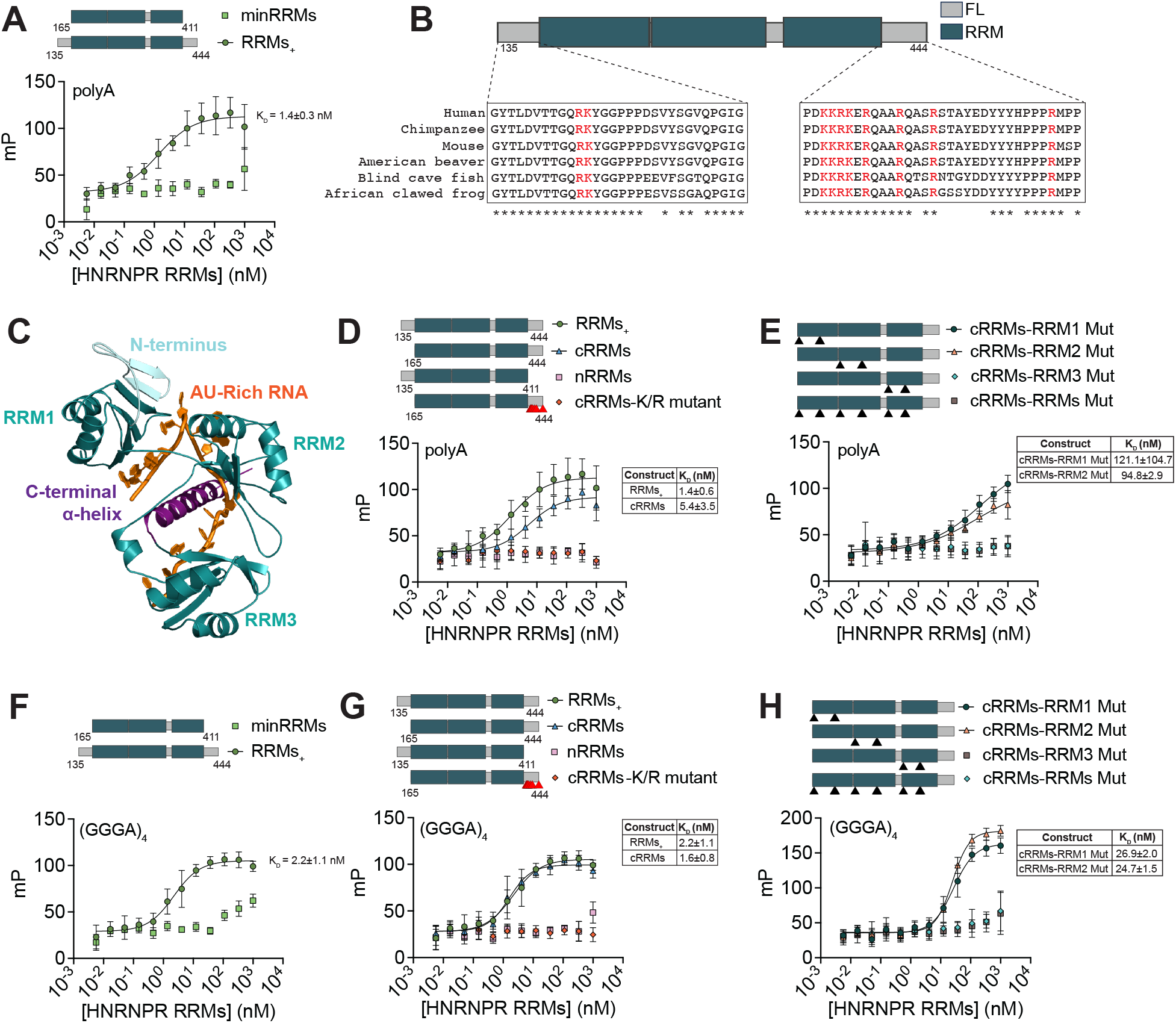
HNRNPR interacts with RNA through an extended RRM domain. A) Top, schematic of HNRNPR minRRMs (Residues 165-411) and RRMs_+_ (Residues 135-444). Bottom, FP binding curves (N=3) for HNRNPR RRMs incubated with polyA RNA. Data are mean ± standard deviation (SD). B) Sequence alignment of HNRNPR RRMs N- and C-terminal regions across species. Stars indicate complete conservation across the selected species. Residues in red are charged. C) AlphaFold3 prediction of HNRNPR RRMs (teal) with AAAUUAAAUU RNA (orange). Residues N-terminal to RRMs are colored in light blue. The predicted α-helix C-terminal to RRM3 is colored in purple. D) Top, schematic of HNRNPR RRM constructs tested. Red triangles represent arginine or lysine to alanine substitutions. Bottom, FP binding curves (N=3) of HNRNPR RRMs incubated with polyA RNA. Data are mean ± SD. E) Top, schematic of HNRNPR RRM mutants. Black triangle indicates a substitution to alanine. Bottom, FP binding curves (N=3) of HNRNPR RRM mutants incubated with polyA RNA. Data are mean ± SD. F) Top, schematic of HNRNPR minRRMs (Residues 165-411) and RRMs_+_ (Residues 135-444). Bottom, FP binding curves (N = 3) for HNRNPR RRMs incubated with (GGGA)_4_ RNA. Data are mean ± SD. G) Top, schematic of HNRNPR RRM constructs tested. Red triangles represent arginine or lysine to alanine substitutions. Bottom, FP binding curves (N=3) of HNRNPR RRMs incubated with (GGGA)_4_ RNA. Data are mean ± SD. (H) Top, schematic of HNRNPR RRM mutants. Black triangle indicates a substitution to alanine. Bottom, FP binding curves (N=3) of HNRNPR RRM mutants incubated with (GGGA_)4_ RNA. Data are mean ± SD.

Previous studies have described additional structural elements that extend beyond the core RRM fold to be important for RRM stability and RNA-binding (*e.g*., PTBP1) (44, 45). Sequence alignments of the HNRNPR RRM flanking sequences across species showed conservation that extends beyond the canonical RRM and include charged residues (Rs and Ks) which can be important for making RNA contacts (**Fig. 4B**). To investigate structural features that may be required for RNA-binding in the flanking regions, we modeled the HNRNPR interaction with AAAUU RNA using AlphaFold 3 (46). This model revealed a predicted α-helix C-terminal to the third RRM that engages RNA (residues 412-444, **Fig. 4C**). The predicted α-helix appears to “clamp” the RNA opposite RRM3 and multiple charged residues within the helix contact the RNA providing a testable model for the role of this α-helix (**Fig. S4A**).

We generated constructs with either the C-terminal or N-terminal flanking regions (cRRMs and nRRMs, respectively) and found that only the C-terminal flanking region conferred binding (**Fig. 4D**), while the N-terminal region was dispensable (**Fig. 4D**). Furthermore, we mutated arginines and lysines within the C-terminal flanking region, which led to a complete loss of AU-rich RNA binding (**Fig. 4D**). Thus, the C-terminal flanking region and the charged residues within this region are critical for RRM interactions with AU-rich RNA.

We next sought to determine which of the RRMs are essential for binding as previous work has shown that not all RBDs within an RBP necessarily interact with RNA, *e.g*., U2AF homology motifs (47). We made alanine substitutions for three aromatic residues within the conserved RNP motifs known to ablate RRM-RNA interactions (48). We tested individual RNP mutants in the context of all three RRMs and the C-terminal flanking region (residues 165-444, cRRMs). Mutation of RRM1 or RRM2 individually retained binding, albeit with decreased affinity (**Fig. 4E**). However, mutation of RRM3 alone completely disrupted binding (**Fig. 4E**). As expected, mutation of all three RRMs together also resulted in loss of RNA binding, indicating that the C-terminal flanking region alone is insufficient for RNA binding and that RRM3 is a critical mediator of AU-rich RNA binding (**Fig. 4E**).

*RNA G-quadruplexes:* To our surprise, we found that the RRMs_+_ also bound rG4 with high affinity, thus the HNRNPR LCD itself was not the only portion of HNRNPR that bound rG4s (**Fig. 4F**). As was the case for AU-rich binding, the minRRMs (no flanking sequences, 165-411) did not bind rG4s (**Fig. 4F**). Similarly, the C-terminal flanking region was essential for the rG4 interaction and only the RRM3 mutations diminished binding (**Fig. 4G, 4H**). These data are somewhat at odds with previously published works showing that disordered regions and not RRMs are the primary interactor with rG4s (21, 49, 50). However, this implies a dynamic relationship between dual recognition of single-stranded and structured RNA-RBP interactions.

### rG4s displace AU-rich sequences for HNRNPR binding

We have identified a portion of HNRNPR (RRM3 with C-terminal flanking region, cRRM) that is critical for binding to completely distinct RNA sequences and RNA structures. These results raise an intriguing question regarding whether HNRNPR can simultaneously bind rG4s and an AU-rich motif. How multiple RNA binding domains within a protein interplay to engage RNA is not well understood. To investigate domain interplay, we performed a series of competition experiments using FL HNRNPR. Competitive FP assays with labeled or unlabeled AU-rich (derived from the top bound *k*mer) or rG4 RNAs (**Fig. 5A, 5B**) were performed. We found that bound AU-rich sequences could be displaced by an unlabeled rG4 (**Fig. 5A**). However, an unlabeled AU-rich sequence was unable to displace an rG4 (**Fig. 5B**). We propose that cRRM are a nexus for rG4 or AU-rich binding, perhaps being a critical site for dictating target selection.

**Figure 5.**
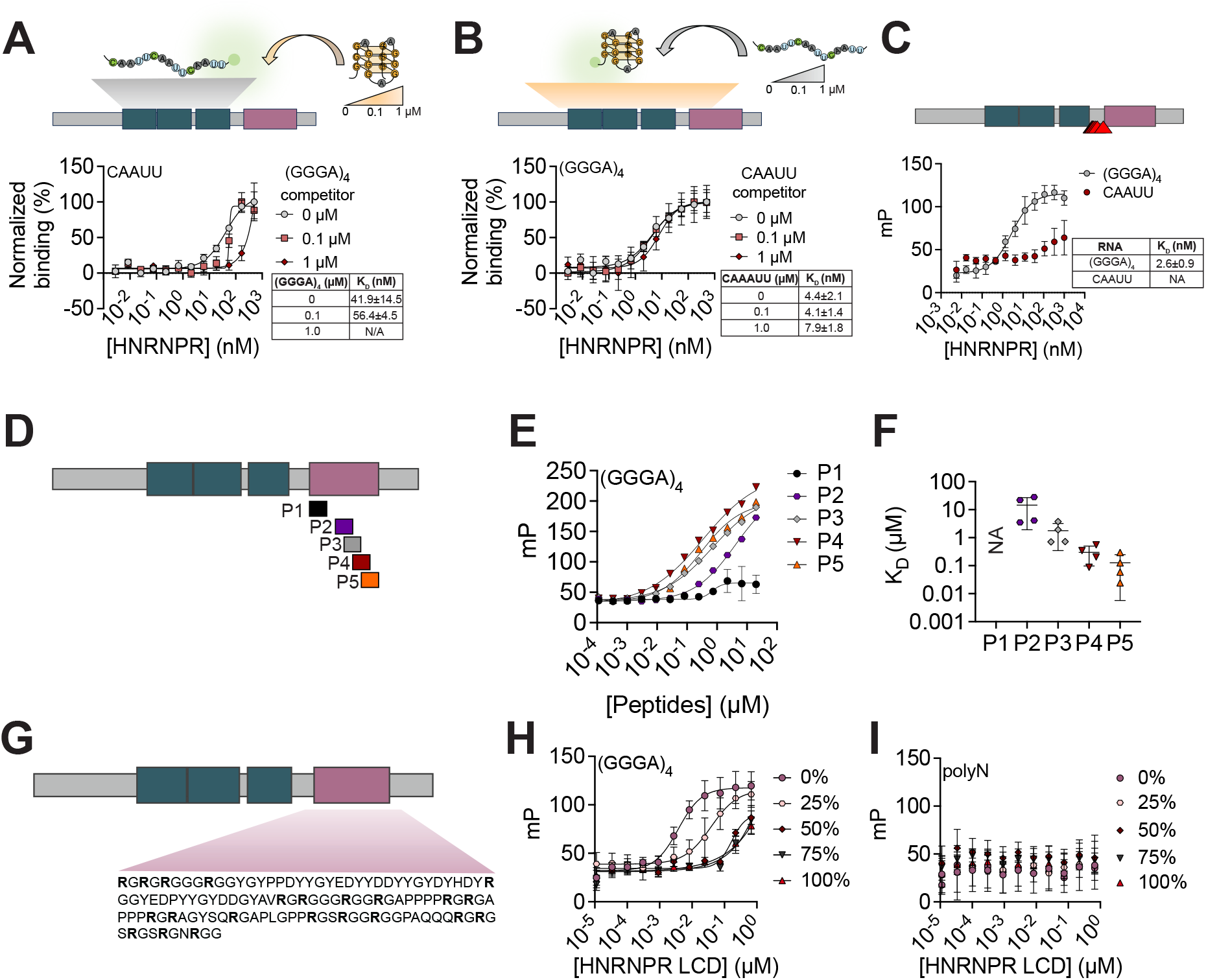
HNRNPR binding duality and dissection of LCD-rG4 binding. A) Top, experimental schematic of the competition FP assay, where FL is incubated with a 6-FAM-labeled AU-rich RNA and an unlabeled rG4 (GGGA)_4_ is added at increasing concentrations. Bottom, FP binding curves (N=3) of the competition FP assay. Data are mean ± standard deviation (SD). B) Top, experimental schematic of the competition FP assay, where FL is incubated with a 6-FAM-labeled rG4 (GGGA)_4_ and an unlabeled AU-rich RNA is added at increasing concentrations. Bottom, FP binding curves (N=3) of the competition FP assay. Data are mean ± SD. C) Top, schematic of FL HNRNPR construct tested. Red triangles indicate substitution for alanine at residues within the C-terminal flanking region. Bottom, FP binding curve (N=3) of FL construct incubated with (GGGA)_4_ RNA and AU-rich RNA. Data are mean ± SD. D) Schematic of HNRNPR highlighting the designed peptides to tile along the LCD. E) FP binding curve (N=3) of HNRNPR LCD peptides incubated with (GGGA)_4_ RNA. Data are mean ± SD. F) Dissociation constant (K_D_) of HNRNPR LCD peptides towards (GGGA)_4_. G) Schematic of HNRNPR highlighting the arginines in the LCD that were substituted by alanines non-sequentially. H) FP binding curves (N=3) for HNRNPR LCD R to A (0% to 100%) mutants incubated with (GGGA)_4_ RNA. Data are mean ± SD. I) FP binding curves (N=3) for HNRNPR LCD R to A (0% to 100%) mutants incubated with polyN RNA. Data are mean ± SD.

Further, because FL HNRNPR also contains an LCD which provided additional rG4 binding capacity but no affinity towards AU-rich RNA (**Fig. 1H**), LCD-rG4 interactions are expected to be insensitive to AU-rich. To test this more directly, we showed that FL HNRNPR bound rG4s even when charged residues in the C-terminal flanking region were mutated (**Fig. 5C**), confirming that HNRNPR has multiple binding sites for rG4s, including some within the LCD. Thus far, we have mapped multiple RNA binding sites with overlapping and distinct specificities within the RBD regions of HNRNPR, highlighting the complexities of RNA binding.

### Multivalent interactions mediate high affinity LCD-rG4 binding

Modes of binding by RRMs have been extensively studied [reviewed in (51)], providing mechanistic details for specific RNA binding. However, how disordered domains mediate nanomolar interactions with RNA is less well understood. To better understand the HNRNPR LCD-rG4 interaction, we assayed multiple peptide fragments by tiling along the LCD region of HNRNPR (**Fig. 5D**). Since binding detection by FP requires a mass-dependent change, and the mass of the LCD fragments is small compared to the rG4 RNA being tested, fragments were synthesized with biotin and complexed with streptavidin for FP. FP results show that fragments bound the rG4 (GGGA)_4_ with varying affinities (**Fig. 5E**), but none displayed binding to randomized RNA (**Fig. S5B**). Notably, none of the fragments displayed affinity comparable to the FL HNRNPR or the LCD; instead, they displayed weaker interactions (**Fig. 5F**). In general, the best binders (P3, P4 and P5, residues 507-567) had multiple RG and RGG repeats, confirming the importance of these short motif repeats in rG4 interactions. We conclude that high affinity rG4 interactions by HNRNPR are mediated by multiple binding sites working cooperatively to engage RNA.

To investigate the importance of arginines more directly, we tested versions of the HNRNPR LCD with arginine to alanine substitutions. Substitutions were generated at random and in increments of 25%, 50%, 75% and 100% R to A (**Fig. 5G, Table S2**). Binding assays showed the best rG4 binder was the WT LCD and mutation of just 25% R to A displayed a 10-fold decrease in affinity (**Fig. 5H**). 50% and 100% R to A versions had greatly diminished RNA binding (**Fig. 5H**). Thus, within the LCD, arginines were the key amino acids for rG4 binding. While the precise number of binding sites will be a challenge to identify, as it is likely they overlap, the observed affinity is best explained by multiple R-containing sites working cooperatively. Altogether, these data demonstrate that HNRNPR, a prototypical RBP, harbors diverse RNA binding capacity involving both unstructured and structured RNA motifs. Additionally, multiple RNA binding sites are found within this protein and together they can dictate affinity and selectivity.

## DISCUSSION

RBPs directly interact with RNA to regulate gene expression. Investigating the mechanism of RNA binding by multi-domain RBPs, like HNRNPR, remains a challenge. Previous studies have shown that individual domains can have unique specificities, overlapping specificity, cooperate to achieve enhanced affinity, or in some cases may not engage with RNA at all (10, 12, 52–55). Our studies indicate that the third RRM of HNRNPR imparts most of the affinity for AU-rich RNA while the first two RRMs appear more dispensable. Recent work has shown that an N-terminal extension of the first RRM of HNRNPR is essential for stability and folding of RRM1 (56). Our recombinant protein constructs of the RRMs (RRMs_+_) did not contain the N-terminal extension of RRM1, raising the possibility that RRM1 was unfolded under our experimental conditions. Despite this, our findings parallel work showing that not all RRMs contribute equally to binding, such as Musashi1/2 wherein one RRM has significantly higher affinity for target RNA than the other (10). We note that while the specificity of HNRNPR FL and the RRMs_+_ was AU-rich, we have not yet investigated the specificity of each RRM individually, and as such non-AU rich motifs may be targets of the individual RRMs. A mechanistic understanding of how all three RRMs work together to bind a natural target RNA will require structural studies.

One unexpected finding was that the minimal HNRNPR RRMs had low affinity for RNA and instead required a charged α-helix C-terminal to the third RRM for high-affinity binding (a fragment not annotated as part of the domain for binding). While C- and N-terminal portions within RRMs have been reported to enhance and diminish RNA affinity [reviewed in (48)], the contribution to binding of regions outside of RRMs is less understood. Previous work found that the RBP, DND1, contains a region N-terminal to its RRM1 that forms an extended surface thought to contribute to high affinity interactions with RNA through specific contacts to the RNA (57). In another case, a region C-terminal to the RRM of PTBP1 acts as a sensor for RNA structure and enhances affinity (44). Intriguingly, this sensor switches from disordered to helical upon RNA binding. In our work, we applied AlphaFold3 to predict RNA binding and observed that the C-terminal charged α-helix next to RRM3 was predicted to make contacts with the AU-rich RNA by “clamping” it into the RRM pocket (**Fig. 4C**). Future structural studies will determine the accuracy of this clamping mechanism. One might envision that regions flanking the RRM may directly contribute to RNA binding or perhaps stabilize the RRM fold itself, an observation that has been made for the first RRM of HNRNPR (56).

Our data also indicate that RRM3 along with the C-terminal helix was able to bind rG4s, an unexpected finding given that most RRM binding appears to be towards unstructured RNAs (51). HNRNPF/H-family proteins have been proposed to bind primarily unstructured G-rich sequences via their RRMs (58, 59); however, some work has indicated their ability to interact with rG4s, particularly in pathogenic repeat expansion RNAs with a propensity to form rG4s (60). HNRNPF/H binding to G-rich RNA has also been shown to destabilize rG4s to modulate splicing outcomes (61). Most known rG4 binders appear to have stretches of intrinsic disorder [reviewed in (62)], which are likely to be the primary drivers of these interactions. RRM3 binding to rG4s not only required the C-terminal helix but also required the charged residues within the helix, consistent with the requirement of arginines for LCD-mediated rG4 interactions demonstrated by us and others (**Fig. 5H**) (20). However, mutating the third RRM to prevent RNA binding also ablated rG4 interactions, even in the presence of the charged helix, indicating a complex binding mode that likely goes beyond a typical RRM-RNA interaction.

The LCD of HNRNPR bound rG4 sequences in multiple assay types and beyond the core fold, we found a preference for strong rG4s with short A- and U-rich loops (**Fig. 3C-H**). Assessment of specificity beyond the core fold was possible because each RNA within our stG4 pool has also been investigated for folding strength in separate experiments, enabling us to demonstrate that rG4 folding strength is important for high affinity binding but features beyond strength are equally important for LCD recognition. While the list of rG4 binding proteins continues to grow, the preferences for specific rG4 features (*e.g*., loops, tetrads, etc.) remains less understood. A limited set of studies have reported RBP specificity for distinct rG4 features (20, 63, 64). For example, HNRNPA1 interacts with rG4s such as *TERRA* via its UP1 domain and RG-rich domain (65). While it has been established that HNRNPA1 can interact with various *TERRA* RNA sequences, only recently has it been shown that the RG-rich domain confers specificity to loop regions of rG4s (64). In a similar example, nucleolin (NCL) has been shown to interact with DNA G4s containing long loops over short loops (63). However, these studies have only assessed preference for a limited set of G4s. We believe the stG4 pool will provide a novel, generalizable framework for dissecting rG4 features that promote RBP-rG4 interactions that can be widely applicable.

A challenge in understanding binding mechanisms of protein-rG4 binding are limited structural studies. X-ray crystallography of FMRP showed that an FMRP-derived RG-rich peptide recognized the stem-quadruplex junction of the *sc1* rG4 (20). Structural studies of DHX36 showed an rG4 interaction via an N-terminal helix that stacks on top of the G-tetrad (66), which perhaps is a similar binding mode to that used by the RRM3-C-terminal helix of HNRNPR. Even within these limited examples of RBP-rG4 interactions, it is clear binding mechanisms are variable.

Consistent with previous work on CIRBP, arginines were critical for high-affinity interactions between HNRNPR and rG4s (49). In the case of CIRBP, analysis of a ∼30 amino acid RG-rich fragment revealed that the arrangement of RG repeats is important for high affinity binding (49). From our own studies, we see that not only are arginines critical, but HNRNPR’s LCD possesses multiple rG4 binding sites that likely cooperate to achieve high-affinity binding. The LCD of HNRNPR is not the only unique domain that contains rG4 binding sites, as the third RRM also harbors such sites, as discussed above. Competition assays with FL reveal that the rG4 can outcompete the AU-rich motif but not the opposite, as the AU-rich motif cannot compete away the rG4 (**Fig. 5A, 5B**). Our interpretation of these data is that the rG4 has unique sites within the LCD, and shared sites with the AU-rich motif with RRM3 and the C-terminal helix. Even at high concentrations of AU-rich competitor RNA, the rG4 is still able to bind the LCD, a portion of the protein that does not appear to interact with the AU-rich RNA. Additional factors as to how AU-rich and rG4 motifs might compete or in some cases co-bind should be considered. These include the steric hinderance of each motif and the impact on folding of the protein that each may potentially have, as rG4s have been shown to improve protein stability [(67) and reviewed in (68)].

Our work highlights the complexity of HNRNPR targets involving both unfolded AU-rich motifs and structured rG4s, each uniquely identified as nanomolar binders. Therefore, the question that remains to be answered is how HNRNPR recognizes these motifs in mRNAs to regulate normal biology. While the present studies lack *in cellulo* binding data, this work serves as a launching point for interpreting targets and binding sites from *in cellulo* analysis.

Our study has several limitations that should be considered. Our work has largely been an *in vitro* analysis, but HNRNPR interactions described herein have not yet been derived *in cellulo* or *in viv*o. This is especially important for rG4 interactions and the complex interplay of AU-rich and rG4 binding. It is likely that additional factors such as the folding strength and dynamics of rG4s will impact potential interactions in the cell. Second, our *in vitro* work requires pure recombinant protein, but as might be expected, highly disordered domains are aggregation-prone and eventually precipitate after purification. To circumvent this issue, we relied on GST-tagged forms of the LCD and RRMs to both limit aggregation and increase the mass of these fragments for FP assays. The known dimerization capabilities of GST should be considered when interpreting binding constants, and at least for the LCD and RRMs, are apparent. We do note that FL HNRNPR was a cleaved untagged form of the protein and largely recapitulated affinity features of the LCD and RRMs. A major strength of our rG4 binding was the use of the stG4 pool, which provides nuanced information on rG4 sequence and structural features that drive binding. Further, HNRNPR binding strength could be correlated to rG4 folding by RT Stop Scores derived using the same pool. However, it is likely that the relationship between rG4 folding and the ability to induce an RT stop is not perfectly correlated, that is, some rG4s may not trigger stops but are still folded. As discussed above and supported by our data, it is plausible that HNRNPR’s LCD can stabilize weak rG4s that do not trigger RT stops.

## METHODS

### In vitro transcription of RNA pools for RBNS

*Random RNA pool (40mer)*: RBNS 40mer DNA pool was synthetized by IDT. *In vitro* transcription was performed using T7 RiboMAX Express Large Scale RNA Production System (Promega, #P1320) following the instruction by the manufacturer.

*stG4 RNA pool*: stG4 pool was designed as previously described (40). Oligos in the stG4 pool were synthesized by Twist Biosciences. Oligo pool was then cloned by polymerase chain reaction and adapted with a T7 promoter sequence. *In vitro* transcription was performed using T7 Ribomax Express Large Scale RNA kit (Promega, #P1320) according to manufacturer protocols for the GTP structural pool. 7-deaza-GTP (Trilink Biotechnologies, N-1044) was substituted in the T7 Ribomax Large Scale RNA Production System (Promega, #P1280) at an equivalent concentration to the GTP for preparation of 7dG RNA pools.

### HNRNPR LCD fragments

Fragments (P1-P5, see sequences in table below) were ordered from and synthetized by the UNC High-Throughput Peptide Synthesis and Array Facility.

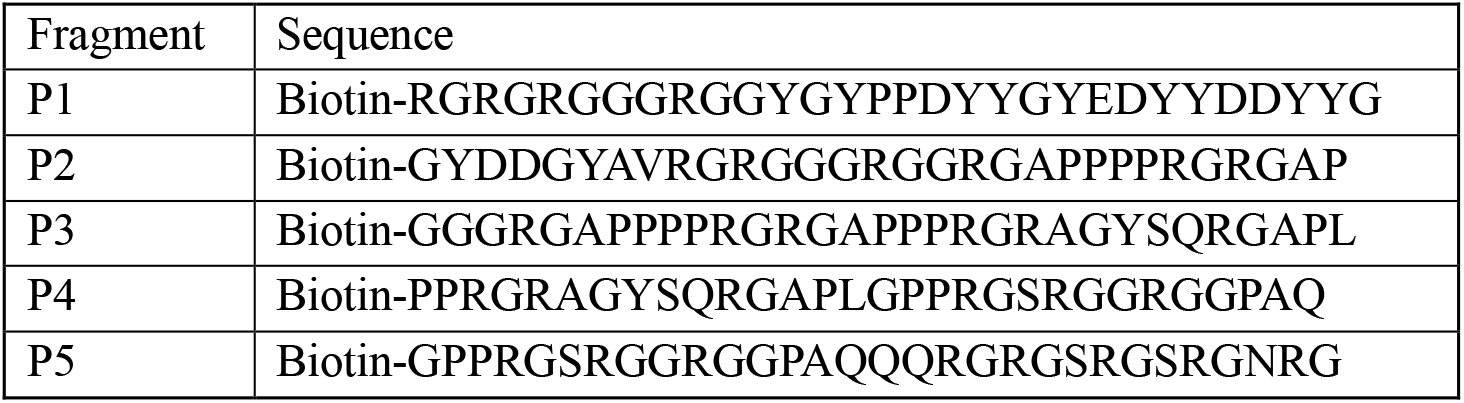

### Protein expression and purification

Plasmids with protein of interest were transformed into Rosetta *E.coli* (Novagen) cells and cultured at 37°C in LB media supplemented with 25 μg/mL chloramphenicol and 100 μg/mL ampicillin to an optical density (OD) of ∼0.8. Upon reaching desired OD, cell cultures were allowed to cool down at 4°C and induced with 0.5 mM Isopropyl β-d-1-thiogalactopyranoside (IPTG) for ∼16 hours at 16°C. Cells pellets were obtained by centrifugation at 4000 x g at 4°C for 13 minutes. Cell pellets were lysed in lysis buffer (1% Triton X-100, 5 mM DTT, 4 mM MgCl_2_, 200 mM NaCl, 20 mM HEPES, 1 tablet of Pierce™ protease inhibitor mini tablet, EDTA-free (Thermo Scientific, #A32955) per 2 liters of culture), sonicated and incubated at 25°C with 250 units of benzonase nuclease (Millipore, #E1014) and 3 units of RQ1 RNase-Free DNase per liter of culture. Cell lysate was then centrifuged at 17,500 x g for 30 minutes and supernatant was collected.

Glutathione (GSH) Agarose (Thermo Scientific, #16101) beads were used for the purification of recombinant proteins. Prior to use, beads were equilibrated with lysis buffer. Cell supernatant was then incubated with beads for 1 hour at 4°C. GST-tagged proteins bound to beads were washed at least 3 times with wash buffer (0.1% triton X-100, 200 mM NaCl, 20 mM HEPES, 3.5 mM EDTA). Proteins were either eluted by elution buffer (20 mM GSH, 50 mM Tris pH 8.0) or with cleavage buffer with PreScission Protease (10% glycerol, 5 mM DTT, 100 mM NaCl, 20 mM HEPES, 0.01% triton X-100, 1 mg/mL in-house PreScission Protease) for ∼1.5 hours at 25°C. If proteins were eluted with GSH, proteins were dialyzed into 150 mM NaCl, 20 mM HEPES. Protein purity was assessed by SDS-Page and Coomassie stain. If sufficient purity was not achieved, proteins were further purified by heparin purification with a gradient from low salt buffer (50 mM NaCl, 50 mM HEPES, 3% glycerol) to high salt buffer (1 M NaCl, 50 mM HEPES, 3% glycerol) on a ÄTKA Pure HPLC and relevant fractions were pooled. Proteins were concentrated using an Amicon Ultra 10 kDa centrifugal filter unit (#UFC8010) and concentration was determined by Pierce BCA Assay Kit (Thermo Scientific, #23208).

### RNA Bind-n-Seq (RBNS)

RBNS was performed as previously reported (33). Briefly, recombinant SBP-tagged proteins were incubated at various concentrations with Dynabeads MyOne Streptavidin T1 (Invitrogen, #65602) for 30 minutes at 4°C in RBNS binding buffer (25 mM Tris-HCl, 150 mM KCl, 3 mM MgCl_2_, 500 mg/mL BSA, 20 units/mL SUPERase IN (Invitrogen, #AM2696)). Following incubation, protein-bead complexes were isolated in a magnetic stand (Invitrogen, #12321D) and unbound proteins were removed. Protein-beads were resuspended in binding buffer and the 40mer RNA random pool was added at a final concentration of 1 mM and incubated for 1 hour at 4°C. After incubation, protein-RNA complexes were isolated in the magnetic stand and thoroughly washed with RBNS wash buffer (25 mM Tris-HCl, 150 mM KCl, 20 units/mL SUPERase IN (Invitrogen, #AM2696)). Subsequently, protein-RNA complexes were eluted twice with RBNS elution buffer (4 mM biotin, 25 mM Tris-HCl) for 30 minutes at 37°C. Eluates were pooled and RNAs were further purified by phenol chloroform extraction. Purified RNAs were heated to 100°C in 150 mM LiCl for 3 minutes and allowed to cool at 25°C for at least 10 minutes. Reverse transcription was performed using RBNS RT primer and SuperScript III Reverse Transcriptase (Invitrogen, #18080044) with a slight modification in the 5X first-strand buffer where KCl was substituted for LiCl. PCR was then performed with RBNS index primers. Libraries were sequenced using an Illumina NextSeq 1000. RBNS was performed in duplicate with two independent protein batches. Enrichment (*R*) was calculated by the frequency of a *k*mer in the protein-bound sample divided by the frequency of same *k*mer in the input pool. Logo generation was performed by selecting the top 15 6mers in R (v. 4.1.0) with R package “rkatss” (v. 0.0.0.9002).

### RBNS with the stG4 pool

RBNS was performed in a similar fashion as described above with some modifications. SBP-HNRNPR FL and SBP-HNRNPR LCD were incubated with Dynabeads MyOne Streptavidin T1 (Invitrogen, #65602) for 30 minutes at 4°C. Following incubation and washing unbound proteins, we either incubated the RBP-beads complex with the stG4 pool in RBNS binding buffer (25 mM Tris-HCl, 150 mM KCl, 3 mM MgCl2, 500 mg/mL BSA, 20 units/mL SUPERase IN (Invitrogen, #AM2696)) or the stG4 pool with 7dG in RBNS LiCl binding buffer (25 mM Tris-HCl, 150 mM LiCl, 3 mM MgCl2, 500 mg/mL BSA, 20 units/mL SUPERase IN (Invitrogen, #AM2696)) for 1 hour at 4°C. Following this step, everything was performed as described above with the only change being that the stG4 pool with 7dG samples were washed with a modified RBNS wash buffer containing 25 mM Tris-HCl, 150 mM LiCl, 20 units/mL SUPERase IN (Invitrogen, #AM2696). Reverse transcription was performed using the RBNS RT primer and SuperScript III Reverse Transcriptase (Invitrogen, #18080044) with a slight modification in the 5X first-strand buffer where KCl was substituted for LiCl. PCR was then performed with RBNS index primers, and libraries were sequenced using an Illumina NextSeq 1000. RBNS was performed in duplicate with two independent protein batches.

Using STAR, raw fastq files were mapped after Illumina adapter trimming, and the reads of every pool sequence per sample were counted. These read counts were then normalized by dividing by the total number of mapped reads per sample, calculating a sequence frequency per sample. Enrichment (*R*) was calculated by the frequency of a sequence (either with guanine or 7dG) in the protein-bound sample divided by the frequency of the same sequence (either with guanine or 7dG) in the input pool. Analysis focused on a subpopulation of the pool derived from synthetic rG4-forming sequences with different sequence characteristics to determine sequence-level drivers of HNRNPR-rG4 binding, described in more detail in (40).

### Fluorescence polarization (FP)

6-FAM fluorescent RNAs were ordered from IDT (**Table S3**). Recombinant proteins were serial diluted with FP binding buffer (150 mM KCl, 20 mM HEPES, 5% glycerol, 0.01% triton X-100, 10 ng/mL Ultra-pure BSA). RNAs were heated to 100°C for 3 minutes in the presence of 150 mM KCl and allowed to cool at 25°C for at least 10 minutes. Serial diluted proteins were then incubated with 5 nM RNA. FP was measured with a CLARIOstar plate reader (BMG Labtech). For each recombinant protein, FP was performed in triplicate with two independent protein batches. For the HNRNPR fragments, FP was performed in triplicate. Binding constants (K_D_) were calculated using the sigmoidal four parameter equation in GraphPad Prism 10.

### FP competition assays

6-FAM fluorescent RNAs and unlabeled RNAs were ordered from IDT (**Table S3**). Recombinant proteins were serial diluted with FP binding buffer (150 mM KCl, 20 mM HEPES, 5% glycerol, 0.01% triton X-100, 10 ng/mL Ultra-pure BSA) and incubated with unlabeled RNAs at the specified concentrations, which were prior heated with 150 mM KCl at 100°C for 3 minutes and rested at 25°C for at least 10 minutes. Likewise, 6-FAM fluorescent RNAs were heated and allowed to cooldown in a similar fashion. Labeled RNAs were then added to the RBP incubated with unlabeled RNA at a final concentration of 5 nM. FP was measured as described above.

### Sequence alignments and structural modeling of HNRNPR RRMs

Sequence alignments of HNRNPR extended RRMs were performed using Clustal Omega (69). Structural modeling was performed using Alphafold 3 (46) with default parameters, using the full length HNRNPR sequence (UniProt Accession: O43390) and the RNA motif: AAAUUAAAUU. Model of RRMs and RNA was visualized with PyMOL (The PyMOL Molecular Graphics System, version 2.1 Schrödinger, LLC.).

### HNRNPR RBNS *k*mer BPP analysis

To study RNA structure and binding, we predicted RNA secondary structure with RNAfold for HNRNPR RBNS data. Based on structural prediction, RNAfold computed base-pairing probabilities (BPP) per nucleotide for each RNA sequence. A *k*mer-sized sliding window was generated along the sequence and calculated the average BPP for each *k*mer within both the randomer RBNS and input data. The average BPP for HNRNPR RBNS were further normalized by the input pool:

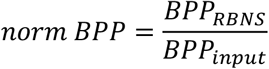

BPP therefore indicated the structural binding bias of each *k*mer. The rank of *k*mers based on the enrichment *R* values and normalized BPP were compared to show the binding preferences of HNRNPR for structure.

## DATA ACCESSIBILITY

Raw FASTQ files and experimental data are available upon request.

## SUPPLEMENTAL DATA

Supplementary data is available online, including Figures S1-S5, Table S1 (rG4 patterns), Table S2 (LCD R to A amino acid information) and Table S3 (RNA oligos used for FP).

## ACKNOWLEDGEMENTS

This work was supported by the U.S. National Institutes of Health (T32GM135095-01 to B.B.G; T32HL069768 to G.A.G.; T32GM148376-01A1 to A.J.; T32HL007149 to J.G.M.; R35GM142864 and R35GM142864-S1 to D.D.), UNC Chapel Hill Simmons Scholar Award (to M.M.A.) and startup funds from UNC Chapel Hill to D.D. Authors would also like to thank Dr. Catherine Eichhorn for her thoughtful feedback on our manuscript.

## AUTHOR CONTRIBUTIONS

**Bryan B. Guzmán:** Writing – Original draft, Writing – Review and editing, Conceptualization, Formal analysis, Investigation, Methodology, Visualization, Data curation. **Grant A. Goda:** Writing – Original draft, Writing – Review and editing, Conceptualization, Formal analysis, Investigation, Methodology. **Alli Jimenez:** Writing – Original draft, Writing – Review and editing, Formal analysis, Investigation, Methodology, Visualization. **Justin G. Martyr:** Writing – Original draft, Writing – Review and editing, Methodology, Visualization, Data curation. **Yue Hu:** Writing – Review and editing, Data curation, Software. **Francisco F. Cavazos Jr**.: Visualization, Data curation, Software. **Maria M. Aleman:** Writing – Original draft, Writing – Review and editing, Supervision, Funding acquisition. **Daniel Dominguez:** Writing – Original draft, Writing – Review and editing, Conceptualization, Investigation, Methodology, Funding acquisition, Supervision, Project administration, Data curation, Resources.

## CONFLICT OF INTEREST

The authors declare no conflicts.

## Notes

### Competing Interest Statement

The authors have declared no competing interest.

