## Supplemental Information for "Contributions of Folded and Disordered Domains to RNA Binding by HNRNPR"

**Table of Contents:**

| | |
|---|---|
| **Figure S1. Additional RBNS characterization** | **S-2** |
| **Figure S2. rG4 pattern reproducibility and FP competition with $(GGGA)_4$ RNA** | **S-3** |
| **Figure S3. Additional binding characterization with stG4 pool** | **S-4,5** |
| **Figure S4. Predicted model of alpha helix interacting with AU-rich RNA** | **S-6** |
| **Figure S5. Additional characterization by FP** | **S-7** |
| **Table S1. Curated list of rG4 patterns used in RBNS** | **.xlsx** |
| **Table S2. LCD R to A mutant amino acid information** | **.xlsx** |
| **Table S3. RNA oligos used for FP** | **.xlsx** |

### Supp Fig. 1

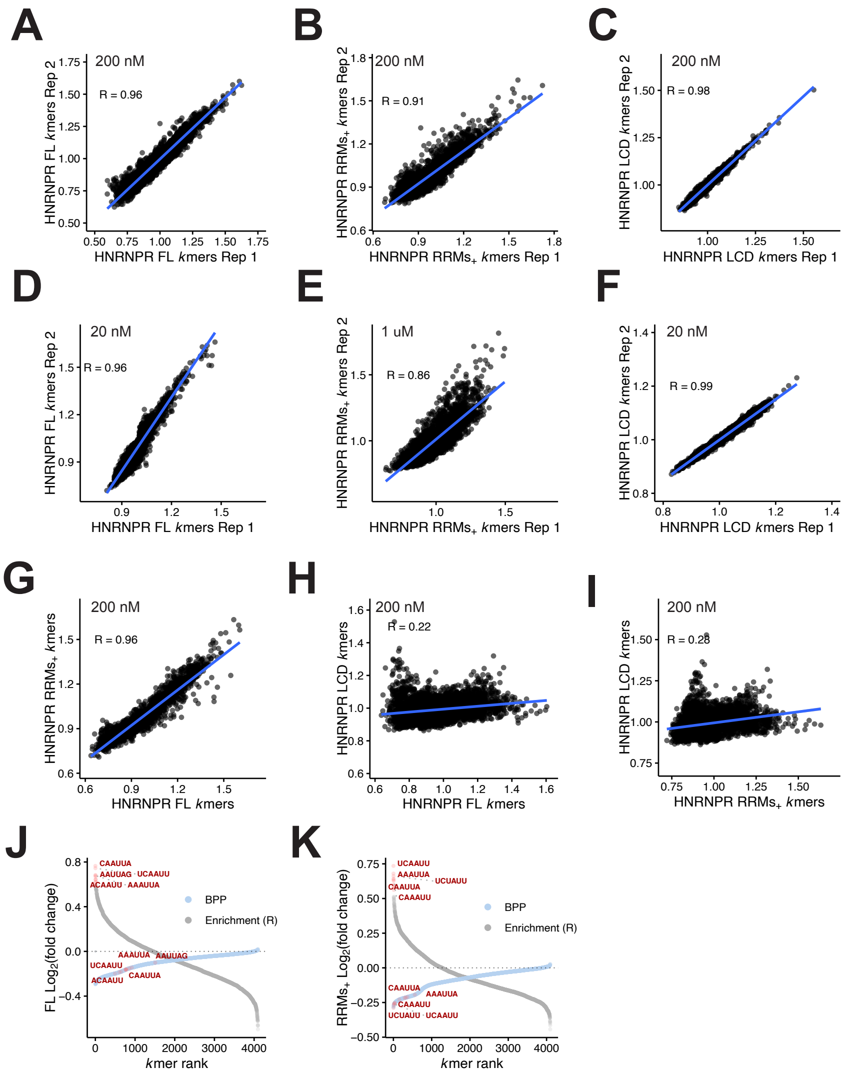


**Figure S1. Additional RBNS characterization.** A) Scatter plot of RBNS enrichment for 6mers of FL Rep 1 (x-axis) versus FL Rep 2 (y-axis) at 200 nM. B) Scatter plot of RBNS enrichment for 6mers of $RRMs_{+}$ Rep 1 (x-axis) versus $RRMs_{+}$ Rep 2 (y-axis) at 200 nM. C) Scatter plot of RBNS enrichment for 6mers of LCD Rep 1 (x-axis) versus LCD Rep 2 (y-axis) at 200 nM. D) Scatter plot of RBNS enrichment for 6mers of FL Rep 1 (x-axis) versus FL Rep 2 (y-axis) at 20 nM. E) Scatter plot of RBNS enrichment for 6mers of $RRMs_{+}$ Rep 1 (x-axis) versus $RRMs_{+}$ Rep 2 (y-axis) at 1 mM. F) Scatter plot of RBNS enrichment for 6mers of LCD Rep 1 (x-axis) versus LCD Rep 2 (y-axis) at 20 nM. Pearson's correlation coefficient is included in each of the plots. G) Scatter plot of RBNS enrichment for 6mers of FL (N=2) (x-axis) versus $RRMs_{+}$ (N=2) (y-axis). H) Scatter plot of RBNS enrichment for 6mers of FL (N=2) (x-axis) versus LCD (N=2) (y-axis) I) Scatter plot of RBNS enrichment for 6mers of $RRMs_{+}$ (N=2) (x-axis) versus LCD (N=2) (y-axis). Pearson's correlation coefficient is included in each of the plots. J) In y-axis, $\log_2$ RBNS enrichment for 6mers of FL (N=2) (grey points) and mean $\log_2$ base pair probability (bpp) of 6mers (light blue points) versus (x-axis) ranked 6mers based on RBNS enrichment. Red points denote top 5 6mers enriched by RBNS. K) In y-axis, $\log_2$ RBNS enrichment for 6mers of $RRMs_{+}$ (N=2) (grey points) and mean $\log_2$ base pair probability (bpp) of 6mers (light blue points) versus (x-axis) ranked 6mers based on RBNS enrichment. Red points denote top 5 6mers enriched by RBNS.

### Supp Fig. 2

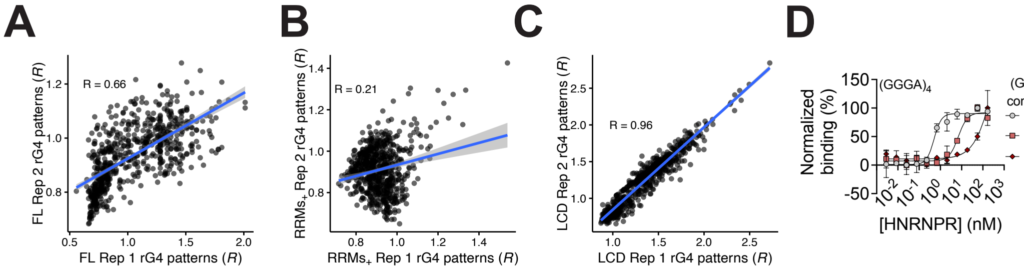


| $(GGGA)_4$ (μM) | $K_D$ (nM) |
|---|---|
| 0 | 1.1±0.7 |
| 0.1 | 14±11 |
| 1.0 | N/A |

**Figure S2. rG4 pattern reproducibility and FP competition with $(GGGA)_4$ RNA.** A) Scatter plot of log2 RBNS enrichment for rG4 patterns for FL Rep 1 (x-axis) versus FL Rep 2 (y-axis). B) Scatter plot of log2 RBNS enrichment for rG4 patterns for $RRMs_+$ Rep 1 (x-axis) versus $RRMs_+$ Rep 2 (y-axis). C) Scatter plot of log2 RBNS enrichment for rG4 patterns for LCD Rep 1 (x-axis) versus LCD Rep 2 (y-axis). Pearson's correlation coefficient is included in each of the plots. D) FP binding curve (N=3) for the incubation of HNRNPR FL with 6-FAM-labeled $(GGGA)_4$ RNA and increasing concentrations of unlabeled $(GGGA)_4$ RNA. Data are mean ± standard deviation (SD).

### Supp Fig. 3

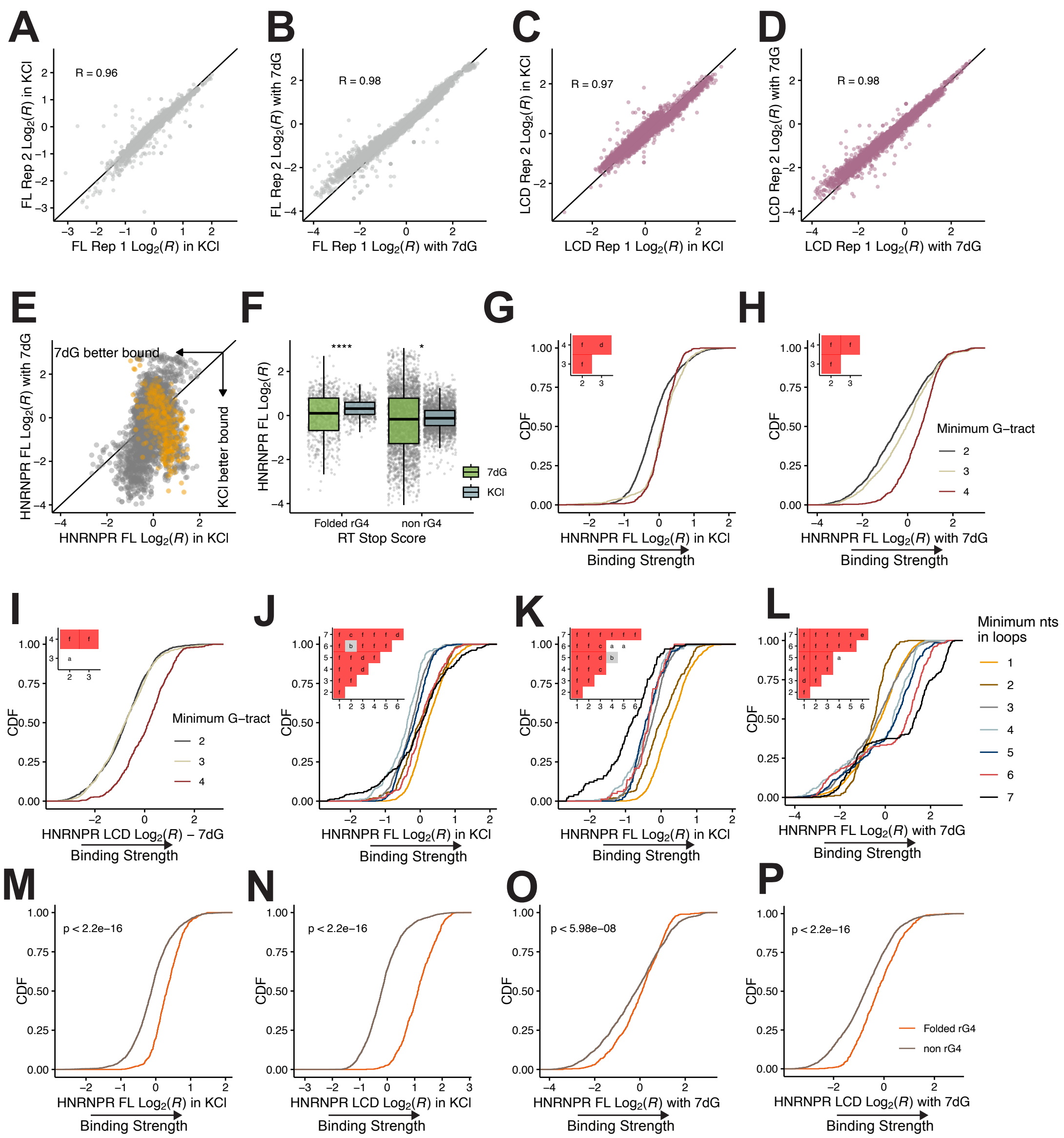

A
B
C
D
R = 0.96
R = 0.98
R = 0.97
R = 0.98
FL Rep 2 Log2(R) in KCl
FL Rep 1 Log2(R) in KCl
FL Rep 2 Log2(R) with 7dG
FL Rep 1 Log2(R) with 7dG
LCD Rep 2 Log2(R) in KCl
LCD Rep 1 Log2(R) in KCl
LCD Rep 2 Log2(R) with 7dG
LCD Rep 1 Log2(R) with 7dG
E
F
G
H
7dG better bound
KCl better bound
HNRNPR FL Log2(R) with 7dG
HNRNPR FL Log2(R) in KCl
****
*
HNRNPR FL Log2(R)
Folded rG4
non rG4
RT Stop Score
7dG
KCl
CDF
HNRNPR FL Log2(R) in KCl
Binding Strength
Minimum G−tract
HNRNPR FL Log2(R) with 7dG
I
J
K
L
HNRNPR LCD Log2(R) − 7dG
Minimum nts
in loops
M
N
O
P
p < 2.2e−16
p < 5.98e−08
HNRNPR LCD Log2(R) in KCl
HNRNPR LCD Log2(R) with 7dG
Folded rG4
non rG4

**Figure S3. Additional binding characterization with stG4 pool** A) Scatter plot of $log_2$ RBNS enrichment of stG4 pool oligos for FL Rep 1 (x-axis) versus FL Rep 2 (y-axis) with guanine in KCl. B) Scatter plot of $log_2$ RBNS enrichment of stG4 pool oligos for FL Rep 1 (x-axis) versus FL Rep 2 (y-axis) with 7dG RNA. C) Scatter plot of $log_2$ RBNS enrichment of stG4 pool oligos for LCD Rep 1 (x-axis) versus LCD Rep 2 (y-axis) with guanine in KCl. D) Scatter plot of $log_2$ RBNS enrichment of stG4 pool oligos for LCD Rep 1 (x-axis) versus LCD Rep 2 (y-axis) with 7dG RNA. Pearson's correlation coefficient is included in each of the plots. E) $Log_2$ RBNS enrichment for FL with 7dG RNA (y-axis) versus $log_2$ RBNS enrichment for FL with guanine in KCl (x-axis). Grey points represent non rG4s (RT Stop Score above -2) and gold points represent folded rG4s (RT Stop Score below -2) F) Boxplot shows the $log_2$ RBNS enrichment for FL for the non rG4 and folded rG4 with 7dG RNA (light green) and with guanine in KCl (light blue). Significance was determined by Wilcoxon test. Significance mark are as follows: * ($p \leq 0.05$), **** ($p \leq 0.0001$). Cumulative distribution function (CDF) of $log_2$ RBNS enrichment for FL separated by the minimum G-tract allowed in an oligo with: G) guanine in KCl and H) 7dG RNA. I) CDF of $log_2$ RBNS enrichment for LCD with 7dG RNA, separated by the minimum G-tract allowed in an oligo. J) CDF of $log_2$ RBNS enrichment for FL separated by the minimum nucleotides in loops with guanine in KCl. K) CDF of $log_2$ RBNS enrichment for FL separated by the minimum nucleotides in loops with guanine in KCl with the removal of long loops (5, 6 and 7 nts) with polyA and/or polyU composition. L) CDF of $log_2$ RBNS enrichment for FL separated by the minimum nucleotides in loops with 7dG RNA. In previous plots, inset shows p-values determined by a two-sided KS test corrected via the BH procedure. Red denotes significance and values are as follows: a (ns), b ($p \leq 0.1$), c ($p \leq 0.05$), d ($p \leq 0.01$), e ($p \leq 0.001$), f ($p \leq 0.0001$). CDF of $log_2$ RBNS enrichment separated by classifying the oligos into folded rG4 (light brown) and non rG4 (dark orange) with guanine in KCl for: M) FL and N) LCD. CDF of $log_2$ RBNS enrichment separated by classifying the oligos into folded rG4 (light brown) and non rG4 (dark orange) with 7dG RNA for: O) FL and P) LCD. In previous plots, p-value was calculated by two-sided KS test.

#### Supp Fig. 4

A

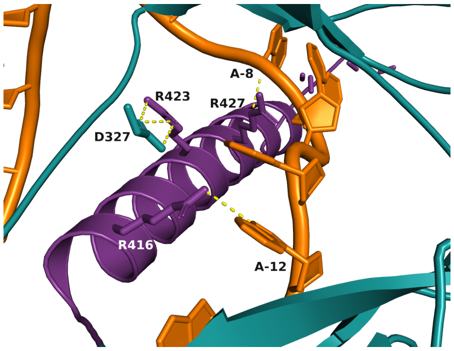


**Figure S4. Predicted model of alpha helix interacting with AU-rich RNA** A) Zoom in view of HNRNPR RRM3 C-terminal α-helix (purple) interacting with AU-rich RNA (orange). Polar contacts between molecules are depicted as a yellow dashed line. Residues engaged in polar contacts are labeled. R416 and R427 in the helix make contacts with bases A-12 and A-8, respectively. R423 makes an intramolecular contact with D327, which is in the loop region between RRM2 and RRM3.

### Supp Fig. 5

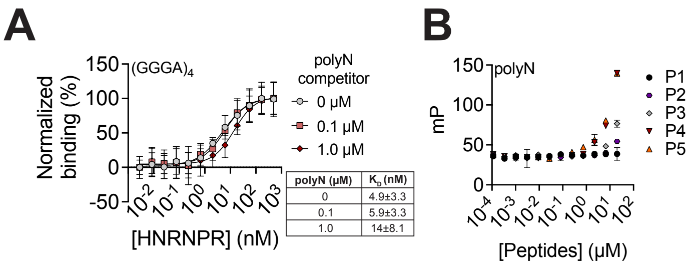


**Figure S5. Additional characterization by FP.** A) FP binding curve (N=3) for the incubation of HNRNPR FL with 6-FAM-labeled $(GGGA)_4$ RNA and unlabeled polyN RNA is added at increasing concentrations. Data are mean ± standard deviation (SD). B) FP binding curve (N=3) of HNRNPR LCD peptides incubated with polyN RNA. Data are mean ± SD.
